# The plant pioneer factor LFY uses trans-kingdom chromatin remodeling mechanisms in vertebrates

**DOI:** 10.1101/2024.12.23.629732

**Authors:** Francisca Ribeiro, Joana Teixeira, Mariana Sottomayor, Jose Bessa

## Abstract

Multicellular complexity is thought to have evolved independently in animals and plants, but the molecular constraints driving this complexity are poorly understood. Genetic networks activated during animal and plant development both involve the regulation of nucleosome occupancy by chromatin remodelers. Pioneer factors, often specific to plants or animals, bind to nucleosome-occupied DNA in a sequence-specific manner, recruiting chromatin remodelers to control lineage-specific genetic networks. In this study, a plant-specific pioneer factor, LFY, when expressed in zebrafish cells, recruits the vertebrate chromatin remodeler Smarca4a to make DNA accessible at LFY binding sites. This allows a plant LFY-controlled promoter, introduced into the zebrafish genome, to become transcriptionally active in a Smarca4a-dependent manner. Our findings suggest that the protein-protein interaction between pioneer factors and chromatin remodelers is a trans-kingdom mechanism likely reflecting ancient eukaryotic events that became a key evolutionary constraint in the path to biological complexity.

## Introduction

Only a few lineages of multicellular life evolved to exceptional levels of complexity, characterized by numerous specialized cell types organized into precise architectures, as seen in animals and plants^1^. This level of organismal complexity implies the presence of intricate genetic programs that require sophisticated mechanisms for controlling genetic information. Different species among the different kingdoms of eukaryotes use chromatin to control the access to DNA information^2^. Pioneer factors are fundamental in this process, having the ability to bind to inaccessible regions of the chromatin and subsequently opening up compacted chromatin structures, making the DNA accessible for other transcriptional regulators to bind and initiate transcription^3^. Consequently, pioneer factors have been shown to activate transcriptional programs that instruct cell fate transitions during development and *in vitro* pluripotency reprogramming^4,5^. Pioneer factors have been functionally studied in several model organisms, the vast majority from the Animalia Kingdom and most of them being representative of vertebrate species^3^. In plants, specifically *Arabidopsis thaliana* (*A. thaliana*), the master floral regulator LEAFY (LFY) has been recently shown to act as a pioneer factor by binding and changing the chromatin state in its target genes^6,7^. LFY, which is lineage-specific to plants^8^, triggers chromatin accessibility of the promoter of the *APETALA1 (AP1*) gene^9^, required for the specification of floral meristem identity, while repressing shoot identity^10^. In order to remodel chromatin, LFY displaces the H1 linker histone after binding to its target DNA sequence in closed chromatin, and recruits the ATPase components of the SWI/SNF chromatin remodelers SPLAYED (SYD) and BRAHMA (BRM), allowing access to closed chromatin in the *AP1* locus^9,11^. The developmental control of DNA accessibility by alterations of the chromatin state seems therefore to be transversal between animals and plants, although complex multicellular development has evolved independently in the two lineages^12^. Here we show that the plant specific pioneer factor LFY^8^, when introduced in zebrafish cells, is able to remodel chromatin by recruiting vertebrate chromatin remodelers, leading to activation of transcription and demonstrating trans-kingdom ultra-conservation of the interaction between chromatin remodelers and lineage-specific pioneer factors.

## Results

### LFY activates a plant target promoter in zebrafish

In *A. thaliana*, LFY triggers chromatin accessibility of the promoter of the AP1 gene^6^. To determine if LFY is able to activate *in vivo* transcription in animals, we first performed transgenesis assays in zebrafish embryos using the promoter of the LFY plant target gene *AP1*, upstream of GFP. After microinjection we observed that the vast majority of embryos showed no GFP expression (Control: 0% with GFP, n=211 vs AP1::GFP: 2.5% with GFP, n=239), while very few embryos showed some GFP positive cells, randomly dispersed (Figure S1A and S1B). These results suggest that the *AP1* plant promoter is mostly inactive in zebrafish cells, having limited predisposition to activate transcription, likely depending on the genomic integration site of the reporter construct. The inactivity of the promoter might be a consequence either of the lack of chromatin accessibility or alternatively because the promoter has no regulatory information to be interpreted by zebrafish transcription factors. To discriminate these two possibilities, we cloned a strong midbrain enhancer (Z48;^13^) downstream of the reporter gene GFP in the *AP1* construct, and we performed a reporter assay as previously. Z48 has the ability to activate transcription in nearby promoters^14^. Therefore, it is reasonable to expect that, if the *AP1* promoter is accessible, it should drive GFP expression in the midbrain of zebrafish embryos. After microinjection, we observed no GFP expression in the midbrain of embryos [Figure S1C and S1D; AP1 Z48: 0% with GFP, n=220 analyzed embryos; Z48 Control (Negative Control): 0% with GFP, n=102; Z48 GATA Control (Positive control): 78.6% with GFP, n=67], suggesting that the *AP1* promoter remains in a closed chromatin configuration in zebrafish cells. To understand if LFY is sufficient to alter the *AP1* promoter into an active state, we microinjected mRNA of *LFY*, at two concentrations (C1:200 ng/µl and C2:100 ng/µl) expected to induce low lethality (Figure S2A), together with the *AP1* reporter construct harboring the Z48 midbrain enhancer (Figure1; Figure S2B and S2C). Upon injection, we observed clear GFP expression in the midbrain of injected embryos, correlating in a dose dependent manner with the used concentration of LFY (Figure1; Figure S2B and S2C; *LFY* mRNA C1, 6.5% with GFP, n=1025 analyzed embryos; *LFY* mRNA C2, 10.2% with GFP, n=974 analyzed embryos; Control mRNA C1: 0% with GFP, n=657; NI control: 0% with GFP, n=680). These results show that LFY is able to activate the *AP1* promoter in zebrafish cells, making it available to interact and respond transcriptionally to the Z48 midbrain enhancer.

### LFY recruits the vertebrate chromatin remodeler Smarca4a to activate a plant target promoter in zebrafish

In *A. thaliana*, LFY needs to recruit the chromatin remodelers SYD and BRM to open chromatin at the LFY binding sites. SYD and BRM have a helicase/SANT-associated (HSA) domain required for the interaction with LFY^11^. Since SYD and BRM are not present in zebrafish, it is reasonable to expect that other vertebrate SWI/SNF chromatin remodelers might interact with LFY. To explore this hypothesis, we compiled all the known SWI/SNF chromatin remodelers from human and zebrafish and performed a domain analysis to locate predicted HSA domains (Figure S3), finding Smarca2 (SMARCA2 in humans) and Smarca4a (SMARCA4 in humans) as top candidates. Additionally, we have aligned the SYD HSA domain with all the vertebrate SWI/SNF chromatin remodelers, finding a match only for Smarca2 and Smarca4a at the exact location of the HSA predicted domain (Figure2A and Figure S3). These results suggest that Smarca2 and Smarca4a might have the potential to bind and be recruited by LFY. If this is the case, the ectopic expression of LFY should generate a dominant negative condition by sequestering Smarca2 and Smarca4a, titrating their binding to other vertebrate pioneer transcription factors and leading to their passive inactivation^3^. For this reason, we have analyzed in detail the phenotypes caused by the ectopic expression of LFY in zebrafish embryos, finding clear phenotypes affecting heart development (heart edema; Figure 2B and 2C). Importantly, the described knockdown of *smarca2*^15^ and the knockout of *smarca4a*^16^ in zebrafish induce heart edema and, additionally, *smarca4a* has been described as important for morphogenesis and patterning of the heart in zebrafish^16^. To confirm the described phenotypes and better compare with the ones obtained in the ectopic expression of LFY, we have knocked down *smarca4a* using a specific morpholino^17^, showing that indeed the knockdown of *smarca4a* partially mimics the phenotypes observed in the ectopic expression of LFY (Figure 2B and 2C). These results suggest that LFY might recruit the vertebrate Smarca4a chromatin remodeler making the *AP1* promoter accessible in the previous reporter assays (Figure 1). To further test this hypothesis, we performed the AP1:GFP:Z48 reporter assay with *LFY* mRNA in the presence and absence of the *smarca4a* morpholino, observing that the GFP expression in the midbrain is suppressed when the *smarca4a* morpholino is present (Figure 3). These results demonstrate that LFY requires the vertebrate Smarca4a to activate the plant *AP1* promoter in zebrafish cells, likely by remodeling chromatin at the LFY binding sites present in the *AP1* promoter.

**Figure 1.**
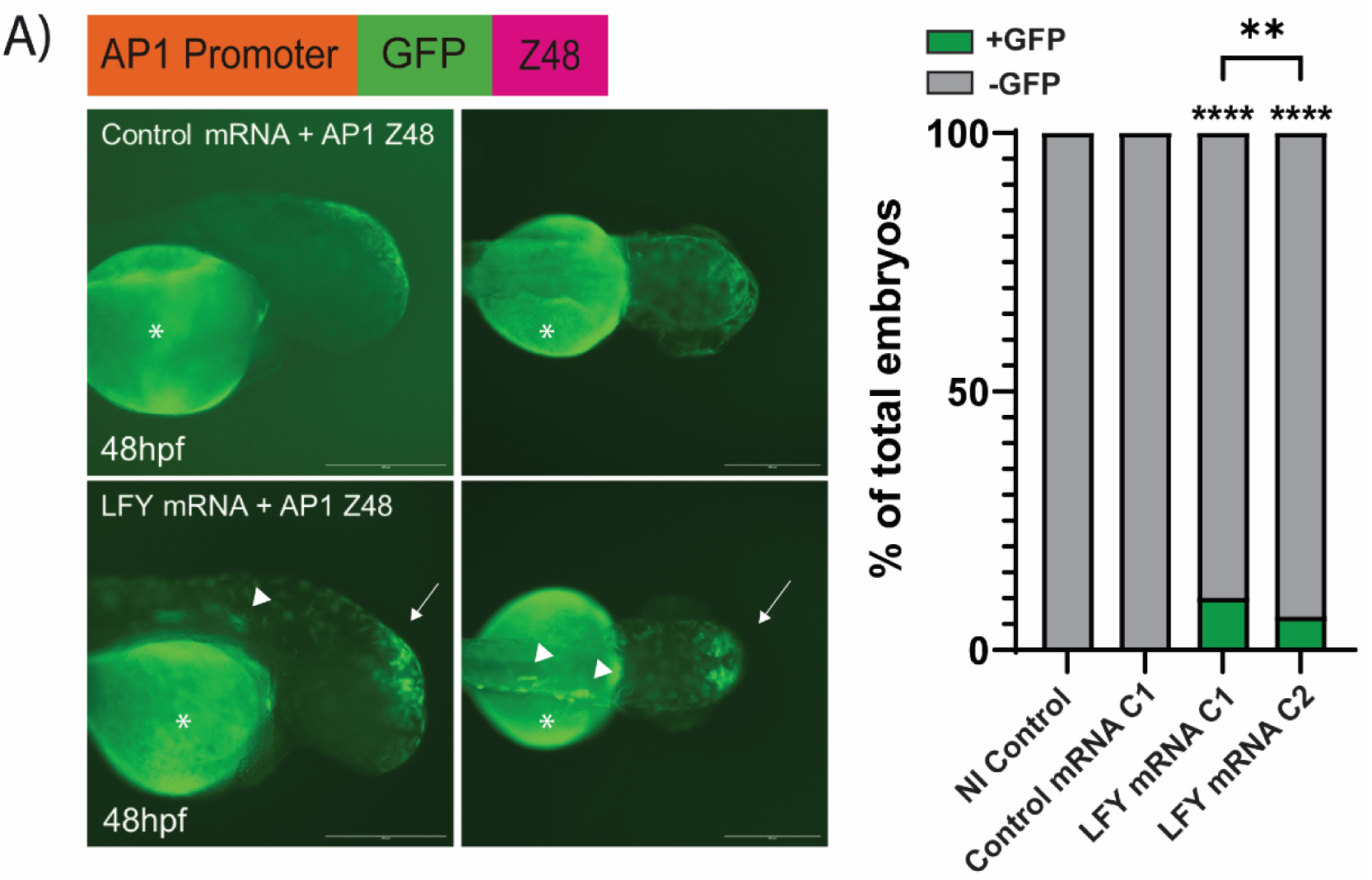
The Arabidopsis thaliana pioneer factor *LFY* can activate *AP1* promoter in zebrafish cells. **(A)** (Left) GFP expression in 48hpf zebrafish embryos co-injected with the AP1 Z48 construct (represented at the top of the figure) and control mRNA (mCherry; 200 ng/μl; top row) or *LFY* mRNA (200 ng/μl; bottom row). Embryos co-injected with the AP1 Z48 construct and *LFY* mRNA show GFP expression in the midbrain (white arrow) and in some muscle fibers (white arrowhead). Asterisk labels autofluorescence in the yolk. Scale bar: 355 μm. (Right) Percentage of zebrafish embryos that show GFP expression in the midbrain in the different conditions. At 48 hpf, non-injected embryos (NI Control, n=680) and embryos co-injected with AP1 Z48 and 200 ng/μl mCherry mRNA (Control mRNA C1, n=657) showed no GFP expression in the midbrain. In contrast, 10.2% of embryos co-injected with AP1 Z48 and 200 ng/μl *LFY* mRNA (LFY mRNA C1, n=974) and 6.5% of those co-injected with AP1 Z48 and 100 ng/μl *LFY* mRNA (LFY mRNA C2, n=1025) showed GFP expression in the midbrain. Percentages were compared using the Chi-square test. Fisher’s exact test was used for sample sizes very small. p- values of less than 0.05 were considered significant (**, p≤0.01; ****, p≤0.0001).

**Figure 2.**
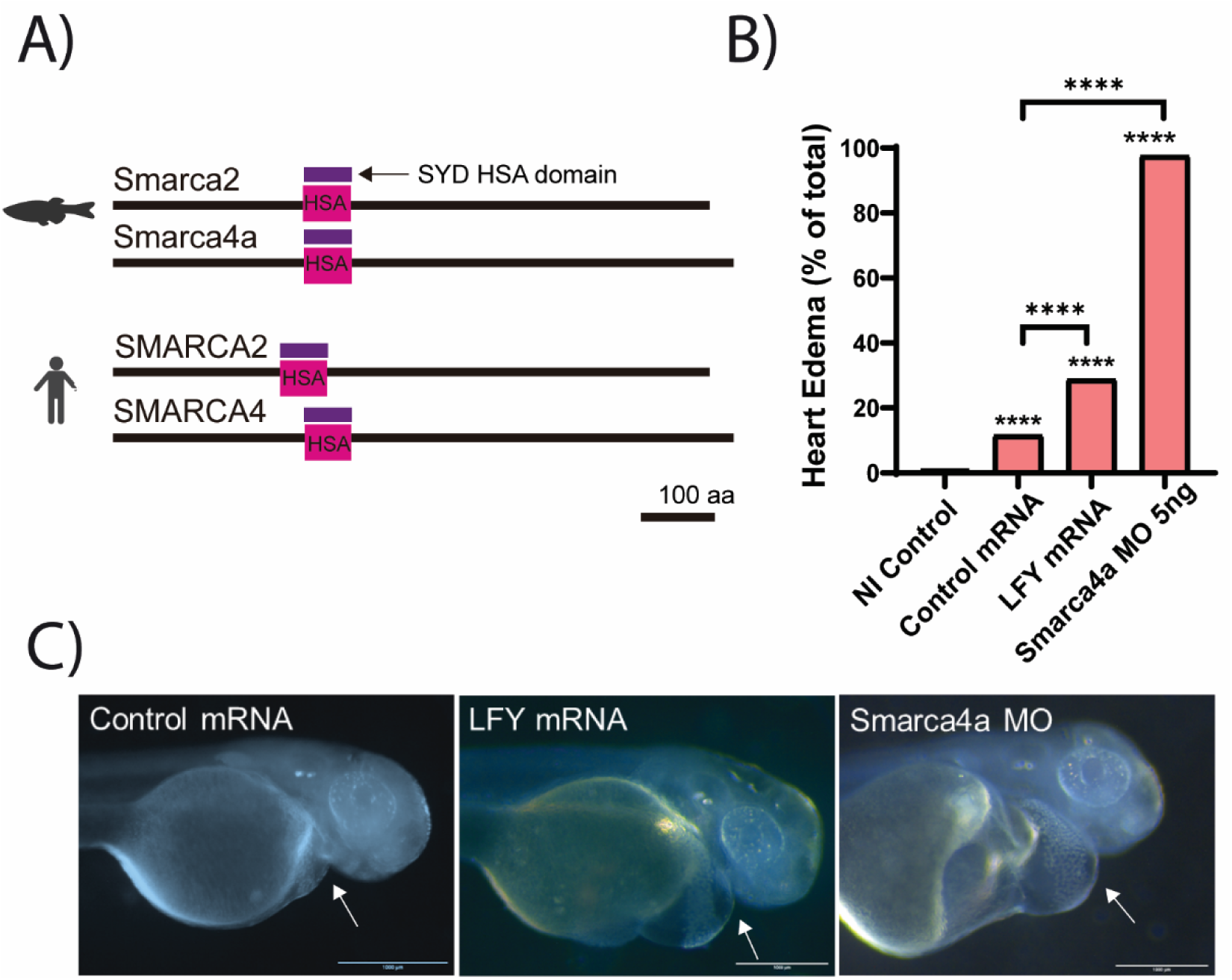
The plant pioneer factor *LFY* might recruit the vertebrate chromatin remodeler Smarca4a in zebrafish cells. **(A)** Graphical representation of the zebrafish (Smarca2, Smarca4A) and human (SMARCA2, SMARCA4) proteins from the SWI/SNF family of chromatin remodelers that contain an HSA domain and respective alignment with the A. thaliana SPLAYED HSA protein domain. Scale is represented by a black bar corresponding to 100 amino acids (aa). **(B)** Percentage of 48hpf zebrafish embryos with heart edema observed when injecting mCherry mRNA (200 ng/μl; control), *LFY* mRNA (200 ng/μl), or Smarca4a morpholino (Smarca4a MO). At 48 hpf, heart edema was observed in 1.1% of the non-injected group (n=820), 11.6% of embryos injected with 200 ng/μl mCherry mRNA (control mRNA; n=449), 28.9% of embryos injected with 200 ng/μl *LFY* mRNA (n=967), and 97.6% of embryos injected with 5ng of Smarca4a MO (n=508). Values were compared by Chi-square and p-values lower than 0.05 were considered significant (****, p≤0.0001). **(C)** Representative images of heart edema phenotypes observed in 48hpf zebrafish embryos microinjected with mCherry mRNA (Control mRNA), *LFY* mRNA or Smarca4a MO. Scale bar: 1000 μm.

**Figure 3.**
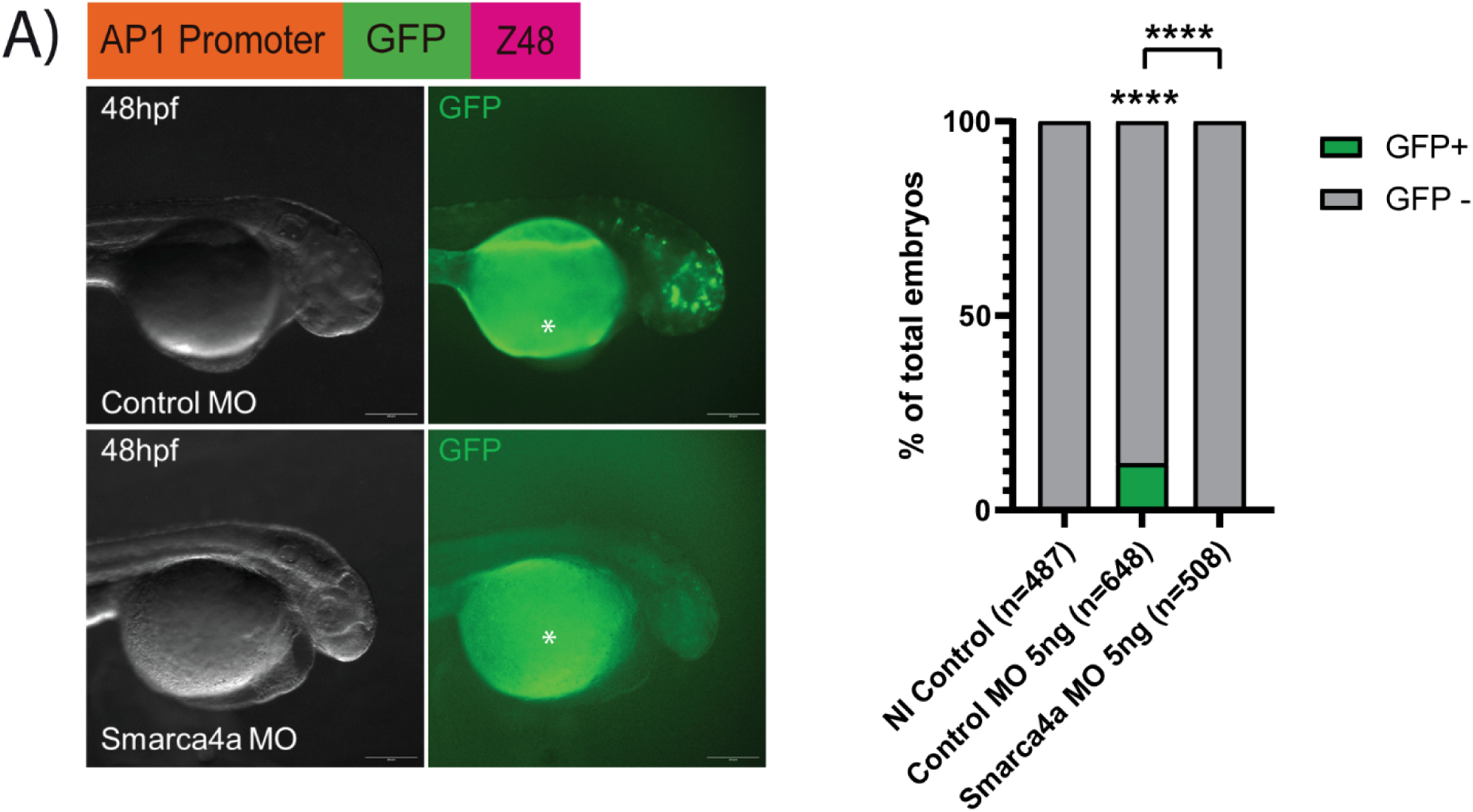
The pioneer factor *LFY* from Arabidopsis thaliana relies on the vertebrate Smarca4a to activate the *AP1* promoter in zebrafish cells. **(A)** (Left) Representative images of GFP expression in 48hpf zebrafish embryos co-injected with *LFY* mRNA, the AP1 Z48 construct and a control morpholino (control MO; top row) or with *LFY* mRNA, the AP1 Z48 construct and the Smarca4a morpholino (Smarca4a MO; bottom row). Autofluorescence observed in the yolk is labeled with an asterisk. Scale bar: 355 μm. (Right) Percentage of 48hpf zebrafish embryos that show GFP expression in the midbrain (green bars) when not injected (NI control), when co-injected with *LFY* mRNA, the AP1 Z48 construct and a control MO or when co-injected with *LFY* mRNA, the AP1 Z48 construct and Smarca4a MO. The non-injected group (n=487) and the embryos co-injected with the Smarca4a MO (n=508) did not show GFP expression in the midbrain, contrasting with the embryos co-injected with the control MO (n=648), showing 12% with GFP expression in the midbrain. Values were compared using Chi-square test or Fisher exact test, depending on sample sizes (****, p≤0.0001).

### LFY changes the genome-wide chromatin accessibility profile in zebrafish

To clarify whether LFY can remodel zebrafish chromatin, we have performed assay for transposase-accessible chromatin with sequencing (ATAC-seq), a method that identifies chromatin accessibility across the genome, in 48 hours post fertilization (hpf) zebrafish embryos microinjected with *LFY* mRNA, and compared the chromatin accessibility profiles with non-injected controls^18^. We have divided the chromatin accessible regions identified in the LFY injected embryos in two groups (Figure S4A): 1) Accessible chromatin regions shared with the control and 2) Accessible chromatin regions exclusively identified in LFY injected embryos. Then we evaluated the percentage of genomic sequences with predicted binding sites of LFY in both of these groups, finding an enrichment in the regions that are exclusive to the microinjected LFY dataset (Figure 4A; Shared: 4.10%; LFY exclusive: 4.60%; p<0.0001, Fisher exact test. Also, Figure S4B). These results suggest that LFY binds to multiple regions of the zebrafish genome containing its binding motif, making these regions accessible. As inferred by the observed phenotypes (Figure 2B and 2C), LFY might be sequestering Smarca2 and Smarca4a to make accessible genomic regions containing LFY binding sites. If so, the presence of transcription factor binding sites (TFBS) motifs for pioneer TFs known to interact with Smarca2 and Smarca4a (Table1 S1) should be less frequent in the LFY dataset of accessible genomic regions than in the control dataset. To test this hypothesis, we have performed a motif discovery of TFBS in open regions of chromatin detected in LFY injected embryos and in control embryos (datasets described in Figure S4A) and we evaluated the difference of the percentage of motifs identified in each of these datasets (Δ%=LFY-Control), for each motif. We observed that pioneer TFBSs known to interact with Smarca2 and Smarca4a are detected less frequently than all the TFBS altogether (Figure 4B, Figure 4C and Table S1). This demonstrates that the ectopic expression of LFY affects negatively the accessibility of the genomic regions where pioneer TFs interacting with Smarca2 and Smarca4a usually bind.

**Figure 4.**
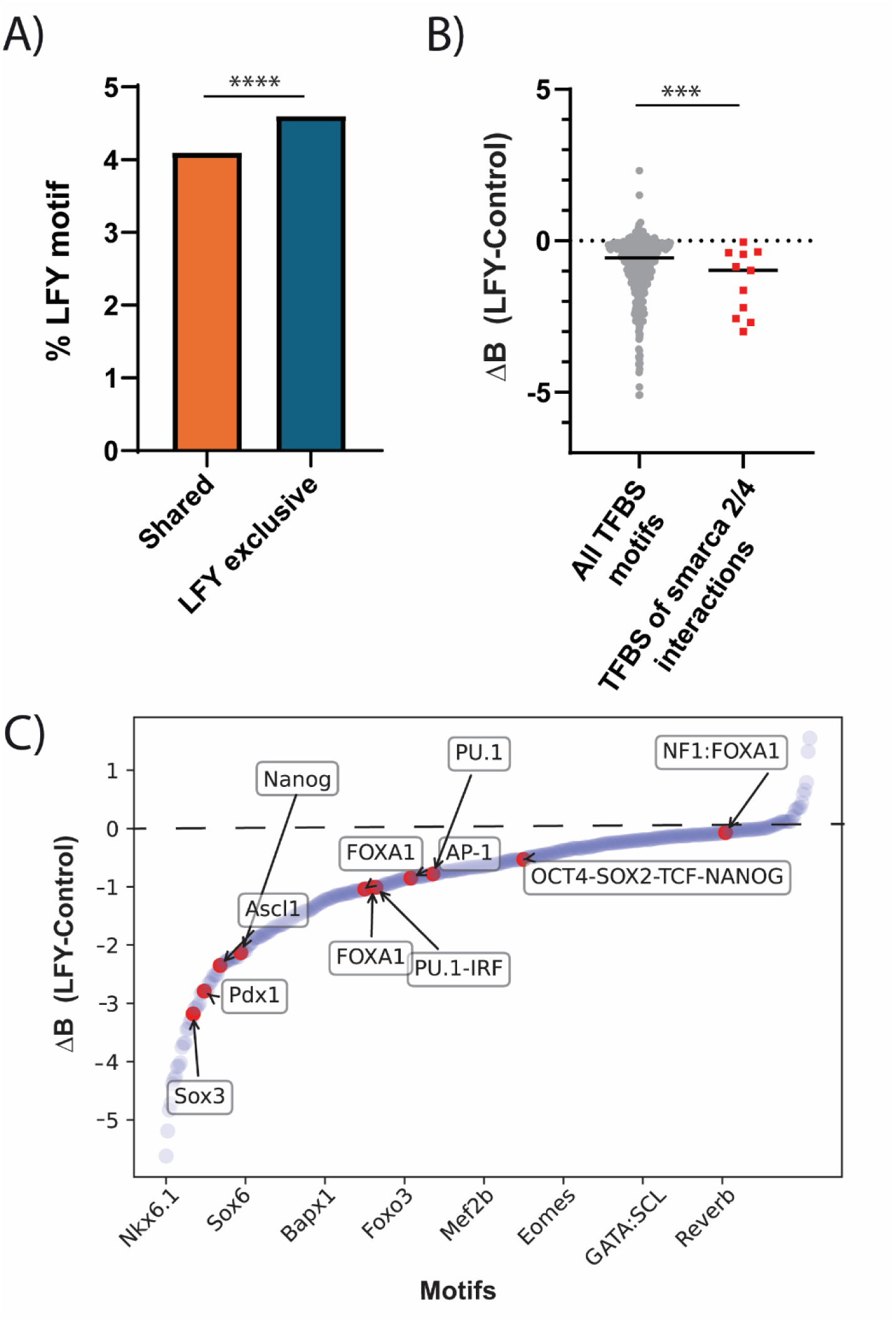
The plant pioneer factor *LFY* reduces the accessibility of genomic regions in zebrafish cells where Smarca2 and Smarca4a interact with vertebrate pioneer TFs. **(A)** Percentage of genomic regions with accessible chromatin detected by ATAC-seq with predicted binding sites of LFY, identified in two groups: The group of accessible regions exclusively detected in LFY injected embryos (blue) and the group of accessible regions shared with the control (orange; see also Fig.S6A; Fisher exact test; ****, p≤0.0001). **(B)** Differential enrichment of TFBS motifs identified in LFY and Control datasets (LFY-Control) for all the motifs (grey dots) comparing to motifs of pioneer TFBSs known to interact with Smarca2 and Smarca4a (list in Table S1; red dots; Fisher exact test; *** p ≤ 0.001). The average is represented by a black horizontal bar. **(C)** Differential enrichment of regions with predicted binding sites of multiple TFs in chromatin accessible regions identified in the LFY injected dataset and in the Control dataset. To determine motif enrichment in each dataset, the difference between the percentage of sequences with a motif and the percentage of background sequences with the same motif was calculated. Positive values represent enrichment in LFY accessible regions, while the negative values represent enrichment in Control accessible regions. Known vertebrate pioneer factors, known to interact with Smarca2 and Smarca4a, are highlighted in red and listed in Table S1.

## Discussion

In *A. thaliana*, LFY recognizes and binds to its cognate motif in nucleosome occupied DNA and triggers chromatin accessibility at the *AP1* promoter, by recruiting the ATPase components of SWI/SNF chromatin remodelers, SPLAYED (SYD) and BRAHMA (BRM)^6,7,19^. We show that an *AP1* promoter reporter is silent in zebrafish embryos, even when cloned together with a strong midbrain enhancer. These results suggest that the *AP1* promoter sequence might contain information that determines a closed chromatin state in zebrafish embryo cells, similar to what happens in *A. thaliana* vegetative organs. Importantly, when LFY is introduced in this reporter background, containing the *AP1* promoter and a strong midbrain enhancer, reporter gene expression is detected in a LFY dose-dependent manner. This indicates that LFY grants access to the *AP1* promoter, becoming a functional target of the zebrafish midbrain enhancer. These results also suggest that the *A. thaliana AP1* promoter includes sequences capable of acting as a minimal promoter in zebrafish. As in *A. thaliana*, the induction of chromatin accessibility by LFY in zebrafish involved the recruitment of a SWI/SNF chromatin remodeler, Smarca4a, one of the closest zebrafish homologs of the *A. thaliana* SYD. This was shown by the significant LFY induction of phenotypes characteristic of Smarca4a depletion, due to Smarca4a’s relocation to the *Ap1* promoter and to other regions of the genome containing the LFY binding motif, and by the suppression of the LFY induced reporter expression when Smarca4a is knocked down. Moreover, assessment of the genome-wide chromatin accessibility upon LFY expression in zebrafish embryos showed a reduction of binding sites of vertebrate pioneer factors known to interact with Smarca4a and an enrichment of regions with predicted LFY binding sites.

Overall, this work presents compelling evidence that LFY, a plant-exclusive pioneer factor that acts as a developmental master switch for flowering, functionally interacts with a vertebrate chromatin remodeler to open chromatin and enable transcription in a vertebrate context. Plant and animal lineages are thought to have diverged at the root of the eukaryotic tree^20^, and independently evolved to reach high multicellular complexity. This complexity is governed by intricate developmental genetic programs that share surprising similarities in the use of master regulators to define temporal and spatial onset of gene expression. However, those similar architectures are controlled by transcriptional regulators that seem to be often nonhomologous^21^. This is the case of LFY, a TF exclusively found in plants, mostly as a single copy gene, that has become a master switch of sexual organ differentiation in flowering plants^8,22^. Remarkably, our work shows that LFY is capable of *in vivo* functional interaction with the vertebrate chromatin remodeler Smarca4a/SMARCA4, namely involved in heart development and in pluripotency reprogramming^5,17,23^. Smarca4a/SMARCA4 belongs to the ancient SWI/SNF family of chromatin remodelers, which are found across multiple Kingdoms of eukaryotes and are therefore thought to have originated in the Last Eukaryotic Common Ancestor (LECA)^24^. This functional protein-protein interaction between a plant-specific pioneer factor that appeared late in the lineage of plants, and an ancient chromatin remodeler originated in LECA, suggests that the interface transcription factor - chromatin remodeler has appeared very early in LECA, likely shaped by the interaction with other ancient pioneer TFs. Indeed, transcription factor families such as Forkhead, SOX, and Homeobox are conserved in animals and plants. Certain vertebrate members of these families, far more functionally explored than their plant counterparts, have been shown to interact with and recruit SWI/SNF remodelers, as in the cases of Foxa1^25^, Sox3^26^ and Nanog^27^, among others. Potential early interactions between pioneer factors and chromatin remodelers at the basis of eukaryotes could have settled a key molecular constraint across the pronounced expansion of new transcription factor classes observed at the root of animal and plant lineages^8^. Searching for the common structural features of the plant/animal pioneer factor domains that interact with chromatin remodelers, and understanding how they originated during evolution, are now questions to be addressed.

Our results underscore protein-protein interactions as an ancestral cornerstone that sustains the evolution of genetic programs responsible for the origin and diversity of multicellular complex organisms. This opens new research avenues that will deepen our fine mechanistic understanding of the regulation and evolution of development, with potential to increase our power to control cell fate for research and applied purposes.

## Methods

### Cloning of the Arabidopsis AP1 promoter in the promoter reporter vectors

Genomic DNA (gDNA) was extracted from a young leaf of an 8 weeks old *A. thaliana* Columbia-0 (Col-0) plant grown under a 16/8h photoperiod, at 22 °C/19 °C, with cool white fluorescent light (70 μmol·m−2·s−1), using 250 mM NaCl, 25 mM EDTA and 0.5% SDS in 200 mM Tris HCl pH 7.5 as extraction buffer followed by standard isopropanol precipitation. Amplification of the AP1 regulatory fragment described in^28^ was performed with i-MAX II DNA Polymerase (iNtRON Biotechnology – R019-220701.51) using the primers 5’ACGAGCTTAGATTCTTTTAGTTTTGC3’ and 5’GAACCAAACAAAACAAAGACCCC3’.

The AP1 promoter was then cloned in the pCR8/GW/TOPO™ TA (Thermofisher) vector, following manufacturer’s instructions, and verified by sequencing. This vector was then used to recombine the AP1 promoter in two Tol2 destination vectors, a Promoter Test Vector with GFP and a Promoter Test Vector with the Z48 enhancer^29^. The first has a GFP reporter gene, a Tol2 transposon, ampicillin resistance and two attB recombination sites located upstream of GFP. The latter has a GFP reporter gene, a Z48 enhancer that drives expression in the zebrafish midbrain when a minimal promoter is cloned upstream of the GFP, a Tol2 transposon, ampicillin resistance and two attB recombination sites located upstream of GFP. The recombination reaction was performed using the Gateway™ LR Clonase™ II Enzyme mix reaction protocol (Thermofisher).

### Cloning and in vitro transcription of the Arabidopsis pioneer TF LFY mRNA

LFY cDNA clones (ABRC DQ447103 and TOPO-U09-C11) were used to amplify LFY using i- MAX II DNA Polymerase (iNtRON Biotechnology – R019-220701.51) and the primers 5’AGATCTATGGATCCTGAAGGTTTCACGAG3’ and 5’GAATTCCTAATCCATGGCGAAACGCAAGTCGTCGCCG3’, with BglII and EcoRI enzyme adaptors. LFY cDNA was then subcloned in pCR8/GW/TOPO™ TA (Thermofisher) vector, following manufacturer’s instructions, and verified by sequencing. LFY cDNA was cloned in the pCS2+ harboring an SP6 promoter for in vitro RNA synthesis^30^. This cloning was done by restriction digestion followed by ligation using BglII and EcoRI and cloned in the compatible EcoRI and BamHI sites located in pCS2+. LFY cDNA cloned in pCS2+ was linearized with XhoI restriction enzyme (Anza, Invitrogen), purified by phenol/chloroform extraction and RNA was in vitro transcribed using SP6 RNA polymerase. Freshly synthesized mRNA was purified using a standard phenol/chloroform purification protocol.

### Zebrafish husbandry and breeding

Zebrafish adults from AB wild-type (WT) strains were maintained in a 14/10 photoperiod (light/dark) in a recirculating housing system, the temperature of the water was set to 26/27°C and the adults were fed three times a day. In order to breed the zebrafishes and obtain embryos, adult male and female fishes (2:3 ratio) were separated the night before spawning into breeding tanks with a partition and maintained at room temperature. In the morning, the fishes were crossed and the embryos were collected after spawning and maintained at 28°C in E3 medium (NaCl, KCl, CaCl.2H2O, MgCl.6H2O and methylene blue), according to standardized protocols (Westerfield, 2000) or in E3 medium supplemented with 0.01% PTU (1-phenyl-2-thiourea) in order to delay pigmentation formation^31^.

### Microinjection of zebrafish embryos and imaging acquisition

One cell stage zebrafish embryos were microinjected in the cell with different mixtures of RNA and DNA, using a 0.58×1.00×100mm microneedle (GB100F-10, SCIENCE PRODUCTS GmbH) previously made in a pooler (PN-31, NARISHIGE). Injection mixtures were prepared as previously described^32^ and mixed a follows: Tol2 DNA plasmids to a final concentration of 25 ng/µl, Tol2 transposase mRNA to a final concentration of 25 ng/µl, LFY or mCherry mRNA to a final concentration of 200 ng/µl or 300 ng/µl and *smarca4a* (5′-CATGGGTGGGTCAGGAGTGGACATC-3′ ; Gene tools ^17^) or control (5′-CCTCTTACCTCAGTTACAATTTATA-3′ ; Gene tools) morpholino to a final concentration of 1ng/nl. Tol2 DNA, mRNA and morpholino were mixed in combination or injected alone. All final solutions had 0.05% phenol red (#P0290, Sigma-Aldrich). Phenotypes and GFP expression were documented at 48 hours post fertilization (hpf) with a stereomicroscope Leica M205FA (Leica Microsystems) on the imaging acquisition system Orca Flash 4.0LT (Hamamatsu Photonics). Image editing was performed with the Fiji software^33^.

### ATAC-seq in zebrafish embryos

ATAC-seq was performed as previously described^34^, with minor changes. One cell stage embryos were injected with mRNA LFY. At 48 hpf, the embryos with heart edema were selected (2 embryos per replicate, with a total of 2 replicates). Following cell lysis, 50000-100000 nuclei were submitted to tagmentation with Nextera DNA Library Preparation Kit (#FC-121-1030, Illumina). ATAC-seq libraries were amplified using KAPA HiFi HotStart PCR Kit (#KK2500, Roche) with the primers Ad1, Ad2.2 and Ad2.3^35^, and further purified with PCR Cleanup Kit (#28104, Qiagen).

### Bioinformatics

#### Predicting vertebrate chromatin remodelers that bind to the plant pioneer TFs in Zebrafish

Protein domain identification was performed using the Expasy ScanProsite tool (https://www.expasy.org/resources/scanprosite;^36^ and protein domain sequences were obtained using Uniprot (www.uniprot.org; The UniProt Consortium, 2023). Protein domain alignments were performed using BLAST^37^.

### ATAC-seq analysis

High quality raw reads from ATAC-seq (FASTQC v.0.11.5;^38^) for the two replicates of 48 hpf embryos injected with mRNA LFY were trimmed for adapter sequences using Skewer (v.0.2.1; ^39^). The generated ATAC-seq data is available at NCBI Gene Expression Omnibus: GSE283875. All libraries were sequenced on Illumina HiSeq 2500 platform and raw reads were mapped to the reference zebrafish genome (GRCz10/danRer10) using Bowtie2 (v.2.2.6) with parameters “-X 2000 and --very-sensitive”^40^. To avoid clonal artefacts, the duplicated mapped reads were removed using Samtools (v.1.9; ^41^). Mapped reads were filtered by the fragment size (≤120 bp) and mapping quality (≥10). To call for enriched regions, MACS2 (v.2.1.0; ^42^) was used with the parameters “--nomodel, --keep-dup 1, --llocal 10000, --extsize 74, --shift –37 and -p 0.07”. For the ATAC-seq analysis, the two replicates were processed independently. Then, we applied the Irreproducible Discovery Rate (IDR, v.2.0.4;^43^) to obtain a confident and reproducible set of peaks. The same pipeline was applied to analyze non-injected 48hpf embryos dataset from^18^; (NCBI Gene Expression Omnibus: GSE156095).

### Transcription factor binding motifs enrichment present in open chromatin regions

The transcription factor binding site (TFBS) predictor program Hypergeometric Optimization of Motif EnRichment (HOMER v.4.11.1;^44^) was used to identify sequence TFs motifs enriched in LFY mRNA injected and Control samples. To determine motif enrichment in each sample, we calculated the difference between the percentage of sequences containing the motif in the sample and the percentage in the background sequences, as outputted by HOMER. We then compared the enrichments across samples by subtracting the enrichment of one sample from another and ranking the results by magnitude.

## Acknowledgments

We would like to thank Marta Duque for sharing the expertise in the preparation of ATAC-seq samples, Ana Eufrasio for helping in zebrafish embryos microinjection and Sara Freitas for sharing Arabidopsis plants and expertise with processing of Arabidopsis material. The laboratory of JB was supported by: “La Caixa” Foundation (under the grant agreement HR21-01212); Portuguese funds through Fundação para a Ciência e a Tecnologia (FCT) in the framework of the project PTDC/BIA-MOL/3834/2021. JB is supported by the FCT grant CEECIND/03482/2018. The laboratory of MS was supported by Project NORTE-01-0246-FEDER-000063, supported by Norte Portugal Regional Operational Programme (NORTE2020), under the PORTUGAL 2020 Partnership Agreement, through the European Regional Development Fund (ERDF), and by the European Union’s Horizon 2020 Research and Innovation Programme under grant agreements Nos 66898 and 857251.

## Author contributions

Conceptualization: JB, MS; Methodology: JB, MS; Investigation: JB, MS, FR, JT; Visualization: FT, JT; Funding acquisition: JB; Project administration: JB; Supervision: JB, MS; Writing – original draft: JB, FR, JT, MS

