## Supplementary Figures for "The plant pioneer factor LFY uses trans-kingdom chromatin remodeling mechanisms in vertebrates"

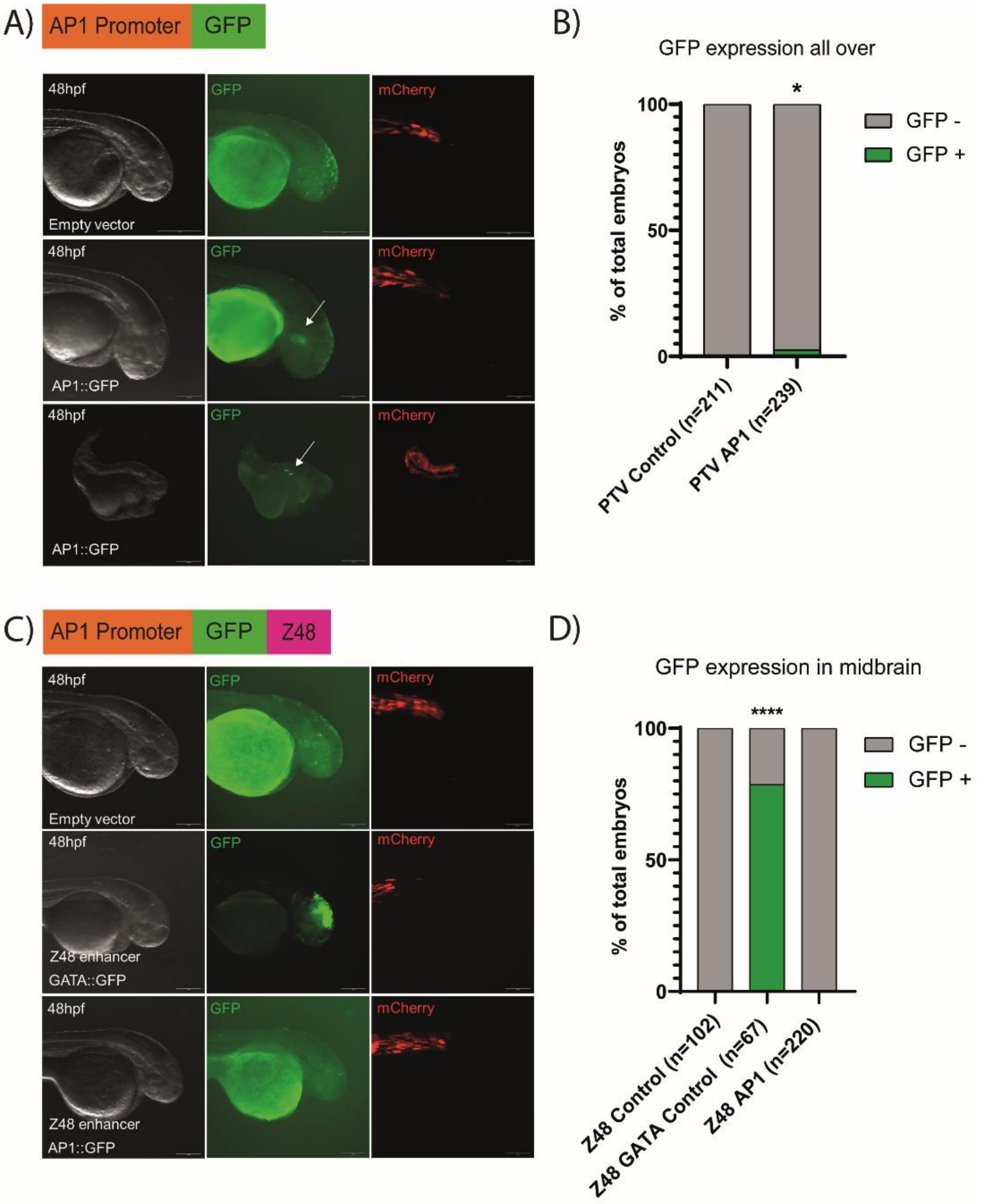


**Figure S1. Assessment of *AP1* plant promoter activity by transgenesis reporter assays in zebrafish cells. (A)** GFP expression in 48hpf zebrafish embryos injected with the empty promoter test vector (PTV) as a control (Empty vector) and the *AP1* construct (AP1::GFP), represented at the top of the figure. All constructs where co-injected with a positive transgenesis control that drives expression of mCherry in muscle cells. Top row: representative image of embryos injected with the empty PTV as a negative control, exhibiting mCherry expression in muscle (right hand image) and showing autofluorescence in the epidermis and yolk (asterisk), but no GFP expression. Middle row: embryo injected with AP1::GFP exhibiting expression of mCherry in muscle (right hand image), showing autofluorescence in the epidermis and yolk (asterisk) and few GFP expressing cells in the head (white arrow; middle image). Bottom row: embryo injected with AP1 exhibiting the mCherry transgenesis control in muscle (right hand image), showing autofluorescence in the epidermis and yolk (asterisk) and some GFP expressing cells in the muscle (white arrow) (middle image); Scale bar: 355 µm. **(B)** Percentage of embryos presenting GFP expressing cells when microinjected with AP1::GFP or control (empty PTV). Control embryos did not show GFP expression. 2.5% of the embryos injected with AP1::GFP show variable GFP expression location. Values expressed as percentages were compared by Fisher exact test. p-values of less than 0.05 were considered significant (*). **(C)** GFP expression in 48hpf zebrafish embryos injected with the empty Z48 PTV (Empty vector) as a negative control, GATA Z48 construct (Z48 enhancer GATA::GFP) as a positive control and the AP1 Z48 construct (Z48 enhancer AP1::GFP), represented at the top of the figure. All constructs were co-injected with a positive transgenesis control that drives expression of mCherry in muscle cells. Top row: Representative embryo injected with the Empty vector, exhibiting the mCherry transgenesis control in muscle (right hand side) and showing autofluorescence in the epidermis and yolk (asterisk) but no GFP expression (middle image). Middle row: embryo injected with the positive control exhibiting the mCherry transgenesis control in muscle (right hand image) and strong GFP expression in the midbrain (middle image). Bottom row: embryo injected with Z48 enhancer AP1::GFP exhibiting the mCherry transgenesis control in muscle (right hand side), showing autofluorescence in the epidermis but no GFP expression in the midbrain (middle image). Scale bar: 355 µm. **(D)** Percentage of embryos presenting GFP expressing cells when microinjected with Z48 enhancer AP1::GFP and controls. Percentage of GFP expression is represented as % of total embryos at 48 hpf. Embryos injected with the Empty vector control did not show GFP expression. 78.6% of the embryos injected with GATA::GFP positive control showed GFP expression. Embryos injected with Z48 enhancer GATA::GFP did not show GFP expression. Values expressed as percentages were compared by Fisher exact test. p-values of less than 0.05 were considered significant.

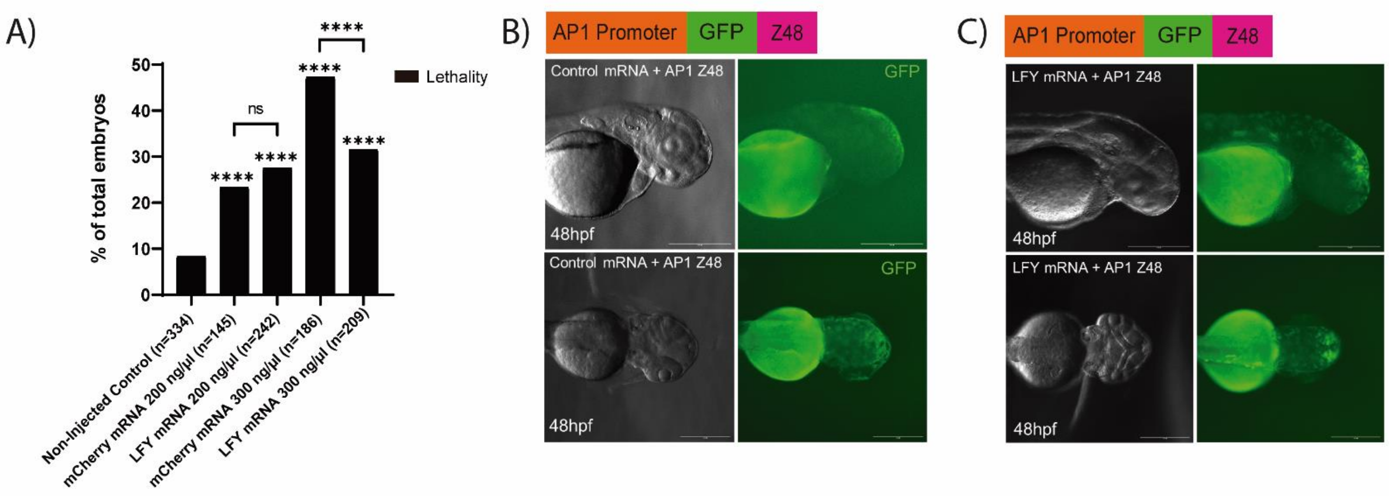


**Figure S2. Assessment of *AP1* plant promoter activity by *LFY* mRNA microinjection in zebrafish cells. (A)** Percentage of dead 48hpf zebrafish embryos upon microinjection of two different concentrations of *LFY* mRNA and mCherry mRNA (control) at two different concentrations, 200 ng/µl and 300 ng/µl. The non-injected group showed 8.4% dead embryos. Embryos injected with 200 ng/µl and 300 ng/µl of mCherry mRNA showed 23.4% and 47.3% dead embryos, respectively. Embryos injected with 200 ng/µl and 300 ng/µl of LFY mRNA showed 27.5% and 31.6% dead embryos, respectively. Values expressed as percentages were compared by the Chi-square test. *p*-values of less than 0.05 were considered significant. **(B)** Representative embryo co-injected with 200 ng/µl of mCherry mRNA and the AP1 Z48 construct, showing autofluorescence in the epidermis and yolk (asterisk) and no GFP expression in the midbrain. Top and bottom rows represent lateral and dorsal views of the embryo, respectively. Scale bar: 355 µm. **(C)** Embryo co-injected with 200 ng/µl (C1) LFY mRNA and the AP1 Z48 construct, showing autofluorescence in the epidermis and yolk (asterisk). GFP expression was observed in the midbrain (white arrow) and in some muscle fibers (white arrowhead). Scale bar: 355 µm.

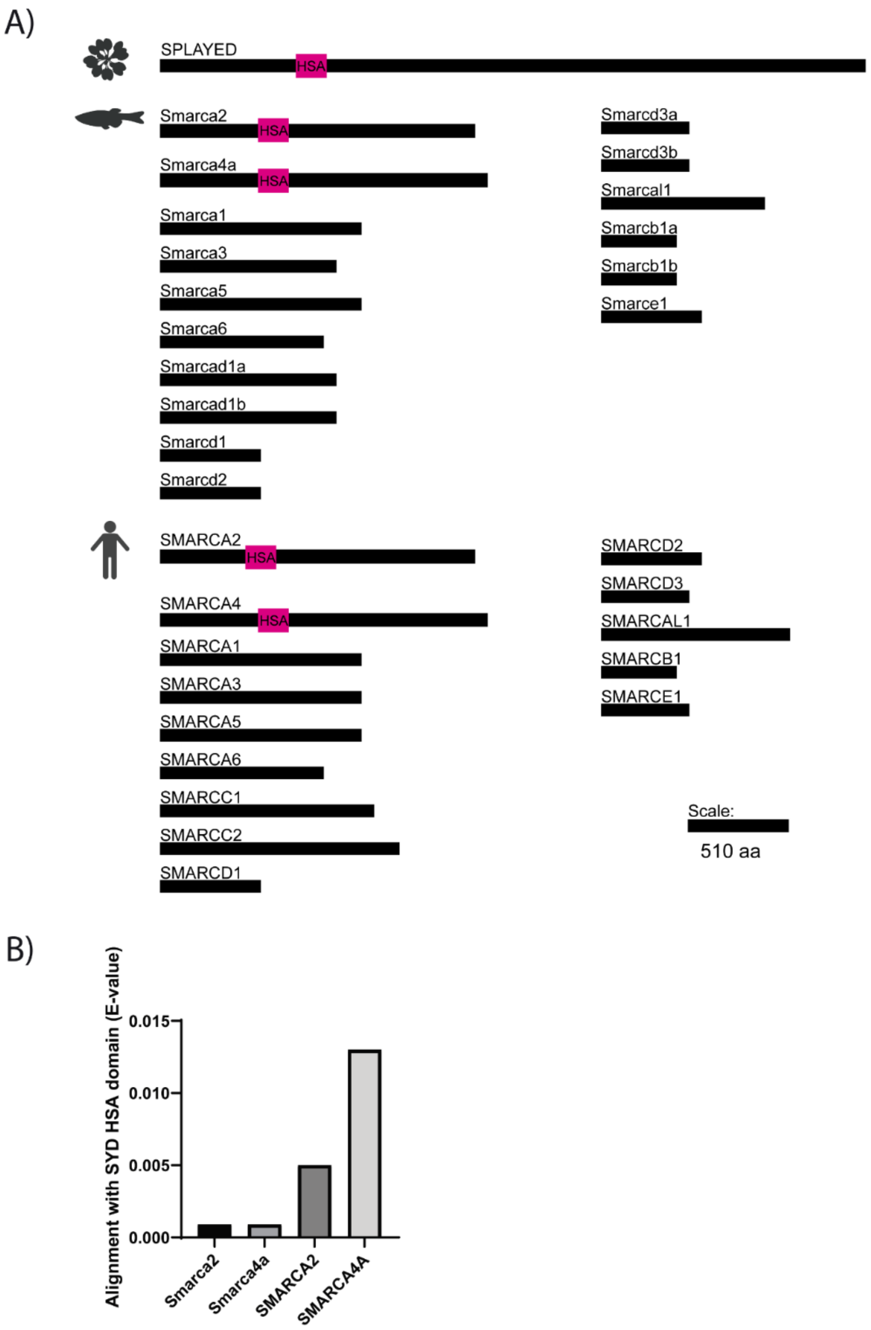


**Figure S3. Graphical representation of the *A. thaliana* SPLAYED protein containing the HSA domain, the Smarca family of proteins and their respective alignment E-values. (A)** Graphical representation of the *A. thaliana* SPLAYED protein containing the HSA domain and the Smarca family proteins, both from zebrafish and human, including the respective HSA domains. Scale is depicted with a black bar corresponding to 510 amino acids (aa). **(B)** Graph showing the alignment E-values, when aligning the SYD HSA domain with Smarca2, Smarca4a, SMARCA2 and SMARCA4A.

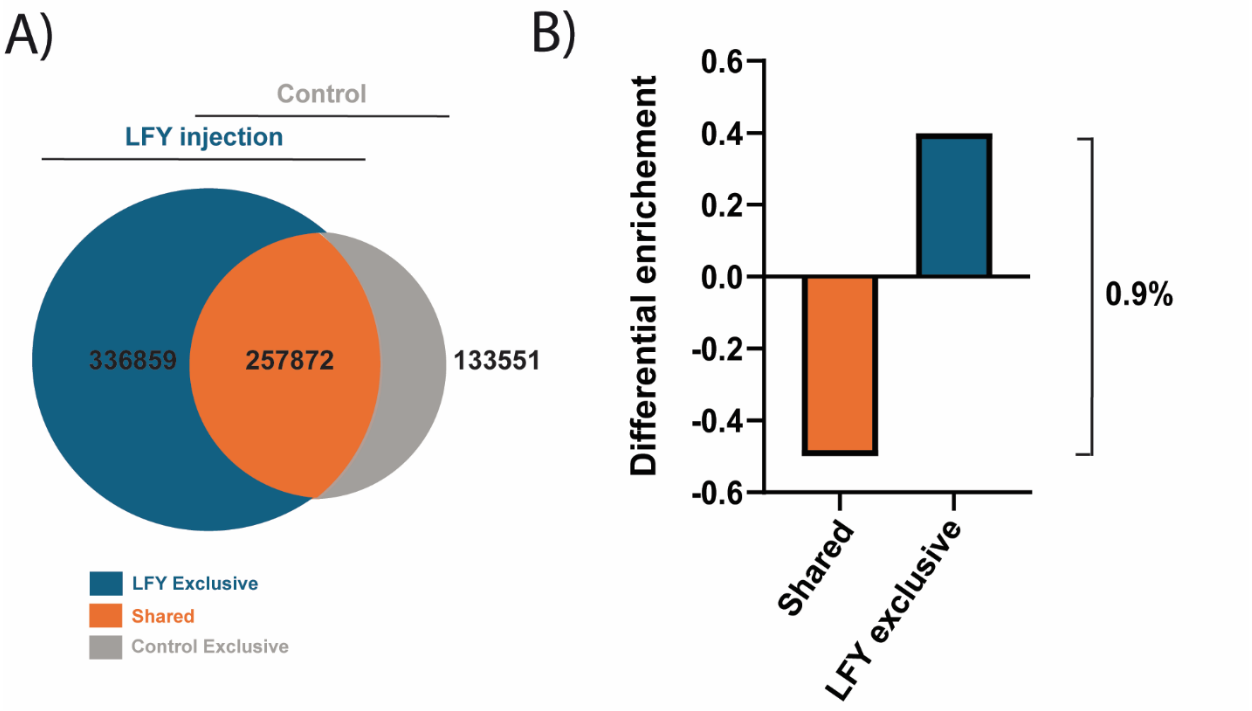


**Figure S4. Chromatin accessibility across the genome, in 48 hpf zebrafish embryos microinjected with *LFY* mRNA and controls and the respective differential enrichment of regions with predicted binding sites of LFY. (A)** Venn diagram showing the overlap of putative open chromatin regions (ATAC-seq) in 48hpf embryos microinjected with *LFY* mRNA (LFY injection; n=594731 regions) and non-injected 48hpf embryos (Control; n=358104 regions). Open chromatin regions exclusive to 48hpf embryos injected with *LFY* mRNA are labeled in blue (LFY Exclusive, n=336859 regions), while the shared open chromatin regions are labeled in orange (Shared, n=257872 regions). The open chromatin regions exclusive to the control embryos are labeled in gray (Control Exclusive, n=133551 regions). **(B)** Differential enrichment of regions with predicted binding sites of LFY in chromatin accessible regions shared with the control (orange) and exclusively identified in LFY dataset (blue). To determine motif enrichment in each dataset, the difference between the percentage of sequences with the motif and the percentage of background sequences with the motif was calculated.
